# Experience from an HTS campaign: Investigation of susceptibility and rescue of SARS-CoV-2 nsp3 protease assay from metal contamination

**DOI:** 10.1101/2025.11.27.690678

**Authors:** Irene Georgiou, Sandra O’Neill, Colin Robinson, Fraser Cunningham, Sean N. O’Byrne, David Gray, Ian H. Gilbert, Duncan E. Scott

## Abstract

Different biochemical assays yield different rates of false positives than others either due to the nature of the enzyme, the technology associated with the assay, or properties of the compounds being screened. Ensuring that the right counter-screens are in place to identify false positives without wasting time and resources on them is of great importance. Herein we describe the results of a high throughput screen (HTS) against non-structural protein 3 (nsp3) protease PL^Pro^, which resulted exclusively in false-positive hits. By triaging hit compounds through purification of metal chelating resin, we identified contamination by either copper or palladium as the most likely source of false positives from the library screening campaign. We then performed a systematic assessment of the vulnerability of nsp3 protease screening to metal contamination and evaluated common additives to combat the inhibitory effects of different metal salts. We further conducted a thorough survey of the literature reports of nsp3 HTS campaigns with a focus on the presence of additives and what metal susceptibility was likely, given the results of our work. We conclude that the majority of reported nsp3 screens are susceptible to copper contamination with a smaller proportion also potentially susceptible to palladium contamination.

## INTRODUCTION

The development of effective therapeutics against SARS-CoV-2 has been of high importance from the start of the COVID-19 pandemic. One of the strategies where vast effort has been placed by the scientific community was exploring druggable targets and identifying suitable small molecule inhibitors.[1-5] Among the most promising targets is papain-like protease (PL^Pro^), which along with main protease (M^Pro^) are responsible for the sequential proteolytic cleavage of the two large polyproteins encoded by SARS-CoV-2, pp1a and pp1ab. PL^Pro^ is one of the functional domains within nsp3, a multi-domain enzyme consisting of 1922 amino acids.[6]

Typical of a cysteine protease, the catalytic triad is situated between the thumb and the palm sub-domain of PL^Pro^, consisting of Cys111, His272 and Asp286 residues. The proteolytic function of PL^Pro^ results in the release of nsp 1-3, which are important proteins in viral transcription and replication.[7] In addition, PL^Pro^ demonstrates de-ubiquitinating and de-ISGylating activities.[8-13] Hydrolysis of the isopeptide bond at the C-terminal of either ubiquitin or ISG15 downregulates the production of cytokines, chemokines and type I interferon, leading to a suppression of the innate immune response. Due to its dual functional role, PL^Pro^ has been an attractive target for drug development. Suitable inhibitors therefore could not only inhibit the viral replication cycle of SARS-CoV-2 but could also prevent the virus evading the host immune system.

The first set of SARS-CoV-2 PL^Pro^ inhibitors reported are the naphthalene based GRL0617 analogues. GRL0617 was originally reported in 2008 as a SARS-CoV-1 inhibitor[14], followed by additional reports exploring the structural activity relationship (SAR) around this pharmacophore.[15-17] With the outbreak of COVID-19, the GRL0617 based inhibitors were repurposed and evaluated by several groups against SARS-CoV-2 PL^Pro^, with GRL0617 demonstrating 2.4 μM inhibitory activity.[9, 18-22] Further SAR optimisation resulted in a new generation of analogues where the naphthalene core was replaced with either 2-phenylthiophene, leading to XR8-23 [23, 24], or with a different aryl motif leading to Jun12682, GZNL-P36 and PF-07957472 with all three demonstrating *in vivo* efficacy.[25-27] Recently two PL^Pro^ inhibitors which are structurally diverse to the GRL-based ones have been reported. This includes an imidazo[4,5-*b*]pyridine-based scaffold and WEHI-P4.[28, 29] In addition to the optimisation of the GRL inhibitors, various other strategies have been applied to identify alternative inhibitors. Among these strategies has been identifying PL^Pro^ inhibitors that would bind covalently to the catalytic Cys111, such as peptidomimetic compounds bearing a reactive α,β-unsaturated warhead or GRL0617-based analogues incorporating a covalent warhead.[30-32] Complementary approaches take advantage of the crystal structure of PL^Pro^ in its apo form and also bound to GRL0617 inhibitors [21, 33, 34] to perform virtual screens using various libraries and different computational methods.[35-41] This led to the identification of naphthoquinone based PL^Pro^ inhibitors.[42] Recent reports suggest the PL^Pro^ inhibitor HL-21 has entered phase I clinical trials but no structural or biochemical information is currently available.[43]

The identification of inhibitors either by a high throughput screen (HTS) or validation of inhibitors resulting from a virtual screen, requires a robust and reproducible primary biochemical assay. One of the most common biochemical assays used in the context of PL^Pro^, uses a FRET-labelled peptide substrate, which bears a native cleavage sequence, recognisable by the enzyme of interest; Dabcyl-FTLRGG/APTKV-5-(Edans).[22, 44, 45] An alternative assay utilises AMC-labelled substrates, (Ub-AMC, ISG15-AMC or RLRGG-AMC, LKGG-AMC); where release of AMC upon proteolytic cleavage results in increased fluorescent signal.[9, 21, 23, 24, 44, 46] Instead of using a small, labelled peptide, an 8.5 kDa ubiquitin moiety attached to Rhodamine 110 has also been utilised as a substrate in a fluorescence assay.[19, 47]

## RESULTS AND DISCUSSION

In an effort to identify novel chemical matter with inhibitory activity against PL^Pro^, which would act as starting points to develop inhibitors with optimised biological and physiochemical properties, we initiated a high throughput screen, **Figure 1**. In this study we employed the SARS-CoV-2 PL^Pro^ fluorescent-based assay that uses an AMC-tagged peptide as a substrate (RLRGG-AMC) and the nsp3 protein (Twinstrep nsp3 (179-1329)-His). An internal library containing 128,794 commercially available compounds was screened against the above-mentioned SARS-CoV-2 PL^Pro^ fluorescent-based assay at a single point (SP) concentration of 30 μM. 1,015 compounds which showed inhibitory activity higher than 48% were retested at a SP concentration. Of those, 393 compounds demonstrated reproducibility in the inhibitory activity and were selected to progress into a 10-point dose response curve (DRC) experiment. The 230 compounds which maintained potency after the DRC study were prioritised by activity and clustered by structural similarity. A set of 37 compounds was progressed into the hit assessment stage, and repurchased or resynthesised solid material of the putative hits was tested in the SARS-CoV-2 PL^Pro^ assay. This resulted in 4 compounds which maintained some level of inhibitory activity, **Figure 2**.

**Figure 1.**
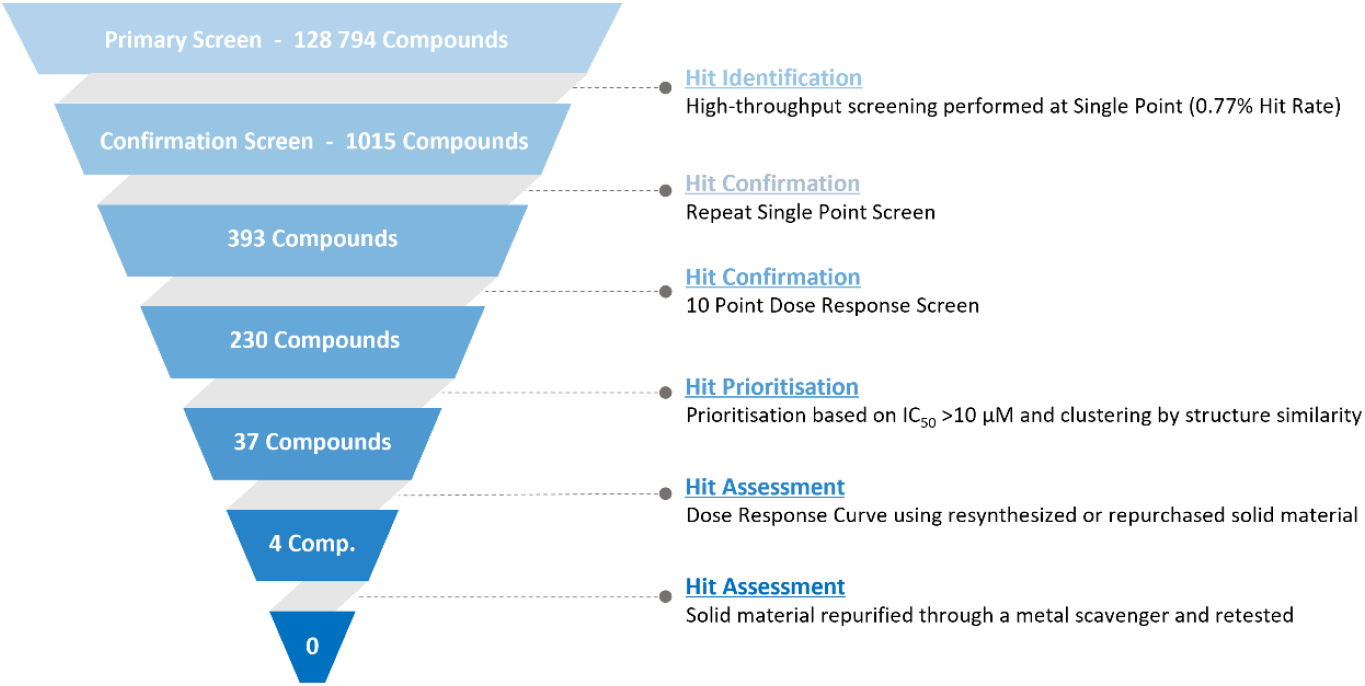
Schematic representation of the High Throughput Screening cascade for the identification of PLP^Pro^ inhibitors.

**Figure 2.**
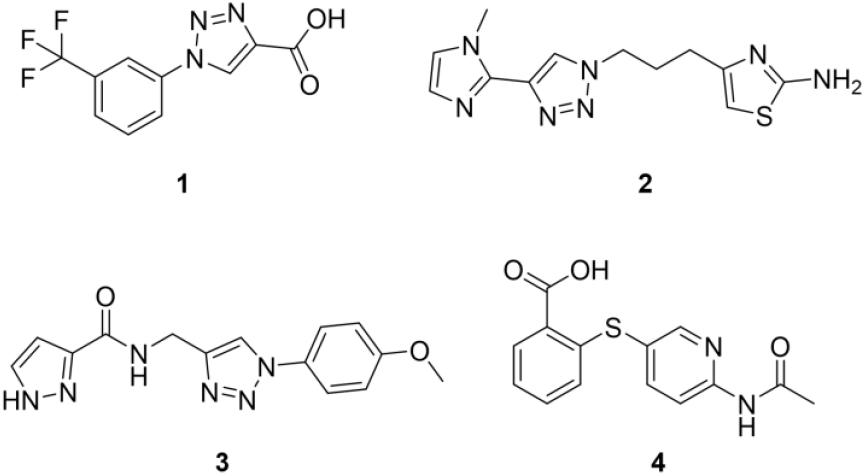
Chemical structures of “putative hits” **1**-**4**.

Among these 4 compounds which in the first instance reconfirmed their potency, 3 compounds had a common structural motif, a 1,4-disubstituted triazole moiety (Compound **1**-**3**; **Figure 2**). The inhibitory activity of these compounds against SARS-CoV-2 nsp3 ranged from IC_50_ 1.3 to 2.6 μM (**Table 1**). The presence of this common triazole structural motif was striking and also noted when performing the structural analysis of the 230 “putative hits”. Hence, this led us to consider two hypotheses. Firstly, that a triazole pharmacophore had been identified and was responsible for the observed biological activity. Or alternatively, a common interference mechanism related to the triazole chemistry might be driving apparent biochemical activity.[48] Taking into consideration that the most applicable synthetic methodology in the formation of substituted triazoles is “click chemistry”, which requires the use of copper-based catalysts, we speculated that potential copper contamination could be a source of potential assay interference. Palladium is ubiquitously useful in chemistry in a very broad range of chemical transformations, and its effects on biochemical activity are also important to understand.

**Table 1.** Measured IC_50_ values of putative hits of various batches against SARS-CoV-2 nsp3.

| Compound | | Purified with metal scavenger resin | IC<sub>50</sub> (μM) | pIC<sub>50</sub> |
| --- | --- | --- | --- | --- |
| **1** | Library solution | | 1.1 | 6.0 ± 0.22 |
| | Commercial solid | | 4.5 | 5.3 ± 0.03 |
|  | Commercial solid | **Yes** | > 100 | < 4.0 |
| **2** | Library solution | | 2.6 | 5.6 ± 0.04 |
| | Resynthesized solid | | 20 | $4.7 \pm 0.13$ |
|  | Resynthesized solid | **Yes** | > 100 | < 4.0 |
| **3** | Library solution | | 1.2 | $5.9 \pm 0.14$ |
| | Commercial solid | | 0.43 | $6.4 \pm 0.27$ |
|  | Commercial solid | **Yes** | > 100 | < 4.0 |
| **4** | Library solution | | 3.7 | $5.4 \pm 0.03$ |
| | Commercial solid | | 2.8 | $5.6 \pm 0.05$ |
|  | Commercial solid | **Yes** | > 100 | < 4.0 |

Part of the assessment of hit quality following the HTS includes retesting of purchased or resynthesized solid material of the 4 hits to confirm activity. Compounds **1, 3** and **4** were commercially available, whereas non-commercial compound **2** was synthesised as outlined in the Supporting Information. In order to exclude the likelihood of any false positives from metal contamination, we implemented an additional purification step for all hit compounds.

One of the most effective ways of removing residual metal impurities from organic compounds is the use of a metal scavenger. Silica or polymer based functionalized metal scavengers are available from various commercial suppliers either as a free powder or in pre-backed columns.[49-51] The different functional groups allow the differentiation in the adsorption levels between metals. In this instance we used Biotage MP-TMT resin, a polystyrene-bound trimercaptotriazine scavenger which can remove various residual metals including Cu and Pd (see Supporting Information).

Repurchased compounds (**1, 3** and **4**) and resynthesised compound (**2**) were tested before and after treatment with a metal scavenging resin for their apparent biochemical activity (**Table 1**). When repurchased compounds **1, 3** and **4** were tested, minor variation in the inhibitory activity were noted as compared to the original activity from the screening library sample (**Table 1**). In the case of compound **1** almost a three-fold drop-in activity was measured. The repurchased batch of compound **3** and **4** showed a comparable IC_50_ to the initial library stock IC_50_. All three compounds were then subjected to the metal scavenger purification, and their activities were reevaluated. Following this additional purification, all compounds showed complete loss of activity (>100 µM). In a similar fashion, resynthesised compound **2** exhibited reduced biochemical potency and complete loss of activity upon metal scavenging. In conclusion, as the biochemical activity of the hit molecules disappeared entirely following metal scavenging, it is apparent that the activity observed initially was most likely driven by palladium or copper impurities. Discrepancies in activity between the resynthesised or repurchased material compared to the original library sample may be down to the levels of contamination. As such, as the nsp3 protein assay appears particularly sensitive to metal contamination, this led us to conduct a systematic study of the effects of assay additives to combat metal-driven false-positives and critically evaluate hits reported more widely in the PL^Pro^ screening literature.

Previous reports have highlighted the problems that inorganic impurities present in screening libraries can cause in the identification and exclusion of false positives in an HTS.[52-54] The standard quality control analysis such as NMR and mass analysis can indicate the presence or absence of organic but not of inorganic contaminants within the samples. Methodologies such as elemental analysis or ICP MS which can determine the presence of inorganic impurities but are not part of routine characterisation techniques, especially in early-stage hit discovery.

Taking into consideration the discrepancy between different batches of compounds and the diminished activity upon purification of the compound through a metal scavenger, we wanted to examine further the effect of metals on the proteolytic function of the protein. Hence, the standard primary biochemical assay was performed in the presence of four different compounds: CuI, Cu(OAc)_2_, Pd(PPh_3_)_4_ and Pd(OAc)_2_ at various concentrations. The concentration of the enzyme (25 nM) and substrate (30 μM) were kept the same as in the original screening assay. All four metal sources showed inhibition of the enzyme with IC_50_ of 0.17 μM, 0.15 μM, 1.8 μM and 2.0 μM respectively (**Figure 2** and **S1**). The same experiments were performed in the presence of additives that could act as a metal chelator or reducer. Specifically, two polyaminocarboxylate-based chelators were used EDTA and EGTA, two thiol-based reagents, DTT and GSH and the phosphopolycarboxylate-based TCEP. All five additives were tested at three different concentrations 0.5 mM, 1 mM and 5 mM.

In the case of Cu(OAc)_2_ the addition of EDTA diminishes the inhibitory activity in all three concentrations (**Figure 3A**). In contrast, the activity is maintained for Pd(OAc)_2_ in the presence of EDTA (**Figure 3D**), suggesting that EDTA alone is not sufficient for removing palladium impurities. The same profiles are noted when EGTA is the additive (See supporting information **Figure S2A** and **S2C**). The addition of DTT to the assay when titrating Cu(OAc)_2_ does reduce the biochemical inhibition due to copper, but in a concentration dependant manner and not entirely at typical assay concentrations (**Figure 3B**). A much cleaner profile was noted for Pd(OAc)_2_ in the presence of DDT (**Figure 3E**), indicating that DDT is efficient at supressing palladium inhibition. A very similar profile to DTT was noted with TCEP as the additive (See supporting information **Figure S2B** and **S2D**). Similar to DTT, GSH showed effective amelioration of palladium-driven enzyme inhibition (**Figure 3F**), but much weaker effects against the inhibitory effects of copper (**Figure 2C**). Comparable outcomes to Cu(OAc)_2_ were noted when the titrant was CuI and to Pd(OAc)_2_ when it was Pd(PPh_3_)_4_ (**Figure S1** and **S3**).

**Figure 3.**
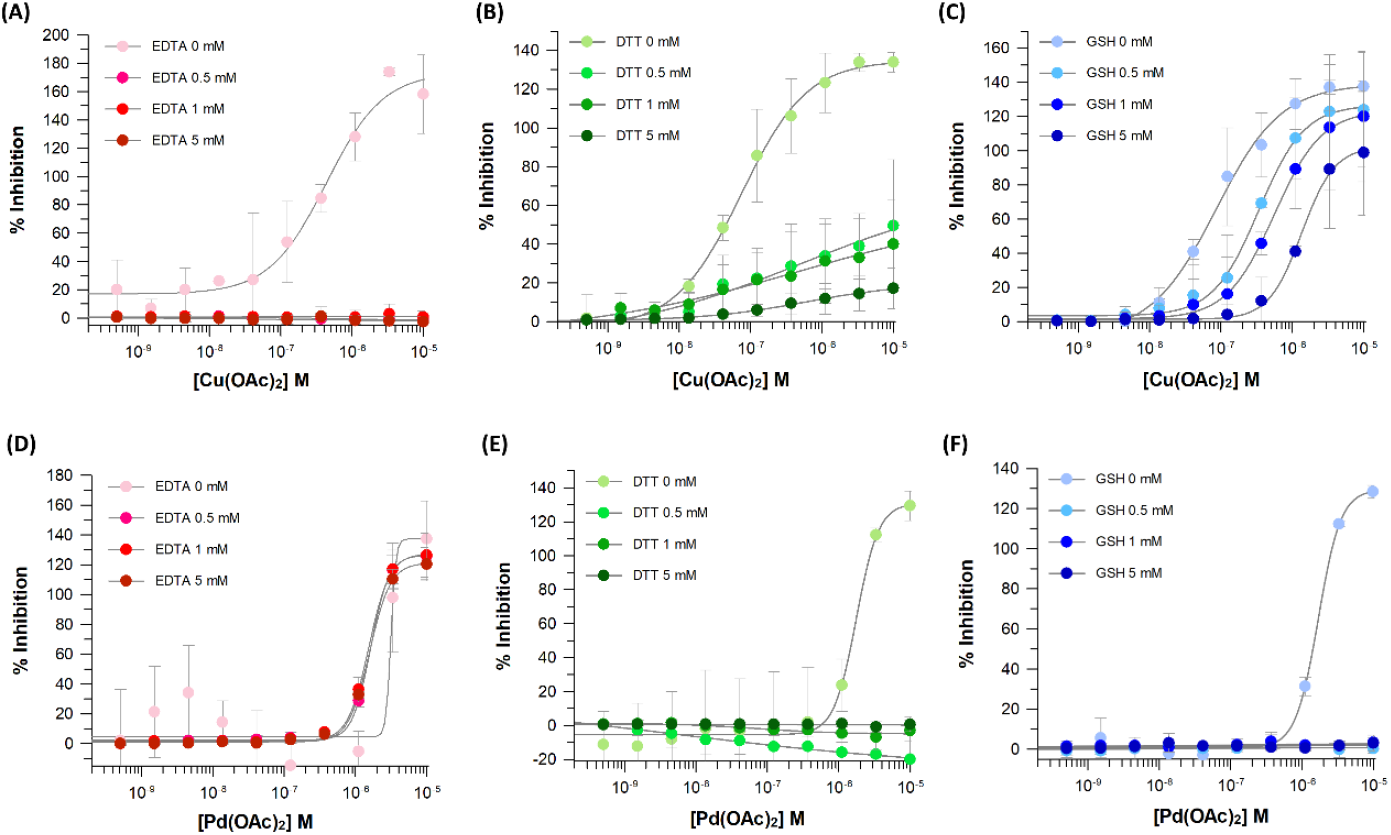
Inhibitory effect of Cu(OAc)_2_ against SARS-CoV-2 nsp3 in the presence of (A) 0.5, 1 and 5 mM of EDTA, (B) 0.5, 1 and 5 mM of DTT, (C) 0.5, 1 and 5 mM of GSH. Inhibitory effect of Pd(OAc)_2._ against SARS-CoV-2 nsp3 in the presence of (D) 0.5, 1 and 5 mM of EDTA, (E) 0.5, 1 and 5 mM of DTT, (F) 0.5, 1 and 5 mM of GSH. Data shown represent mean ± SD from three independent experiments.

On the basis of these findings, a subset of the 37 compounds prioritised from our HTS campaign were rescreened, **Table 2**. Specifically, in addition to 3 of the compounds already outlined (compounds **1**-**3, Table 2**), 12 more compounds (compounds **5**-**16, Table 2**) with diverse chemotypes and a GRL-based positive control (compound **16, Table 2**) were selected. A sample from the library stock for each compound was screened in our SARS-CoV-2 nsp3 assay in both the absence and presence of an additive. The two additives selected for further evaluation were EDTA and DTT. For most compounds the previously reported inhibitory activity was lost in the presence of EDTA or DTT with the exception of the GRL-based positive control, which was not affected by any of the additives. Compound **5** maintained its potency in the presence of EDTA but not with DTT, suggesting a copper impurity was present in the tested sample. For compounds **10, 13** and **15** a significant drop-in activity was observed with DTT, and complete loss of activity with EDTA present. These observations suggest that the number of false positives attributed to metal contamination identified from our SARS-CoV-2 nsp3 assay can be reduced by the addition of a metal chelator(s) in the biochemical assay. For our HTS hits, EDTA did a better job than DTT at removing false positives.

**Table 2.** Measured IC_50_ values of putative hits of various batches against SARS-CoV-2 nsp3 in the absence of a metal chelator and in the presence of EDTA or DTT.

| Compound | Additive |  |  |  |  |  |
| --- | --- | --- | --- | --- | --- | --- |
|  | - |  | EDTA |  | DTT |  |
|  | IC<sub>50</sub> (μM) | pIC<sub>50</sub> | IC<sub>50</sub> (μM) | pIC<sub>50</sub> | IC<sub>50</sub> (μM) | pIC<sub>50</sub> |
| 1 | 1.6 | 5.8 ± 0.18 | > 100 | < 4.0 | > 100 | < 4.0 |
| 2 | 2.5 | 5.6 ± 0 | > 100 | < 4.0 | > 100 | < 4.0 |
| 3 | 1.3 | 5.9 ± 0.19 | > 100 | < 4.0 | > 100 | < 4.0 |
| 5 | 6.0 | 5.2 ± 0.42 | 4.8 | 5.3 ± 0.14 | > 100 | < 4.0 |
| 6 | 1.5 | 5.8 ± 0.32 | > 100 | < 4.0 | > 100 | < 4.0 |
| 7 | 1.5 | 5.8 ± 0.10 | > 100 | < 4.0 | > 100 | < 4.0 |
| 8 | 4.2 | 5.4 ± 0.06 | > 100 | < 4.0 | > 100 | < 4.0 |
| 9 | 2.3 | 5.6 ± 0.27 | > 100 | < 4.0 | > 100 | < 4.0 |
| 10 | 2.0 | 5.7 ± 0.15 | > 100 | < 4.0 | 49 | 4.3 ± 0.15 |
| 11 | 3.2 | 5.5 ± 0.24 | > 100 | < 4.0 | > 100 | < 4.0 |
| 12 | 4.0 | 5.4 ± 0.17 | > 100 | < 4.0 | > 100 | < 4.0 |
| 13 | 0.35 | 6.5 ± 0.01 | > 100 | < 4.0 | 33 | 4.5 ± 0.04 |
| 14 | 1.0 | 6.0 ± 0.15 | > 100 | < 4.0 | > 100 | < 4.0 |
| 15 | 0.92 | 6.0 ± 0.41 | > 100 | < 4.0 | 25 | 4.6 ± 0.06 |
| 16 | 2.7 | 5.6 ± 0.16 | 2.8 | 5.6 ± 0.16 | 3.0 | 5.5 ± 0.12 |

Although the incorporation of an additive in an HTS could reduce the false-positive signal attributed to inorganic impurities, it cannot guarantee the complete removal. As demonstrated by our experiments, the identification or not of a false positive is heavily dependent on the presence of a suitable additive in the biochemical assay, and the type of inorganic contaminant present. With these findings in mind, we reviewed the nsp3 screening literature, with a focus on reviewing the additives used in the SARS-CoV-2 PL^Pro^ biochemical assays reported from 2020 onwards, comprising 28 separate studies (See Supporting Information). The analysis revealed that DTT was used in 65% of reported screens, TCEP in 20% and EDTA in 15% of them. According to our observations, EDTA can diminish the inhibitory activity of Cu-based but not Pd-based contaminants as effectively. Conversely, DTT can prevent inhibition from Pd-based contaminants but not Cu-based contaminants cleanly. The combination of EDTA and DTT as buffer additives therefore might be expected to offer superior protection from both Pd and Cu impurities. A closer look into the reported literature reveals that only two out of the 28 reports use an EDTA and DTT combination.[26, 41] Applying our observations on the differential metal chelation properties of different additives has led us to conclude that 32% of reported PL^Pro^ screening studies are susceptible to Pd contamination and 86% (24 of 28 reports) are susceptible to Cu contamination. Interestingly, Wang *et al*. demonstrated that literature reported SARS-CoV-2 PL^Pro^ inhibitors drastically reduced enzymatic activity compared to the originally reported values and failed to exhibit the same outcome in an orthogonal assay.[45] The authors attribute the inconsistency either to the use of different substrates in the biochemical assay or the possibility of an interference mechanism. In both reports, DTT was the only additive present in the biochemical assay, hence the possibility of Cu being the source of disparity can not be ruled out.

Given our HTS campaign experience and this analysis, we suggest that a close look at screening conditions with nsp3 is advised. Any hits followed up should be re-tested in the presence of chelating additives and ideally with material repurified by metal chelators. Finally, although we have shown that metal contamination of library compounds is one possible source of false positives for nsp3 assays, we also observe that many hits in the literature appear likely to interfere with the fluorescence-based assays, which are typically utilised for this protein, or that the hits reported contain potential redox liabilities or general PAINS flags, for which further controls must be put in place.[55]

## CONCLUSIONS

Common assay interference mechanisms include nonspecific binding, redox compounds, aggregators, interference with the assay readout (i.e. fluorescence) and organic or inorganic impurities.[48] One of the strategies in identifying and excluding compounds that show false positive activity is to include a counter screen as part of the screening workflow. For example, in the case of fluorescence-interfering compounds, examination of the raw fluorescent signal may be possible as is changing the protein assay readout to a non-fluorescence-based readout. False positives which are driven by metal contamination are a little more insidious, as changing the assay format is unlikely to change the compounds’ inhibitory profile. A well-documented approach to overcome this is the incorporation of a suitable additive, either a metal chelator or reducing agent into the assay. This is not always a feasible solution, since the stability or function of the protein maybe be prone to the addition of them. Moreover, a single additive may not remove all relevant interfering metal impurities. In the case of nsp3 we have demonstrated the vulnerability of the protein to both copper and palladium contamination and highly recommend that both EDTA and DTT are included in screening buffers for this target in the future. Ensuring that the compounds becoming part of screening libraries are free of contaminants, is also critical. This is usually difficult to control and document with compounds purchased from commercial sources. Even with in-house resynthesised compounds, it is not common chemistry practise to purify compounds with metal chelators. The case therefore for including metal chelators in the screening buffer is further reinforced.

During the validation of a putative hit from an HTS, regardless of whether the solid material planned to be used is from a commercial source or resynthesized, both can be susceptible to various levels of contamination. As well as an assay designed to minimise false positives, implementation of a purification technique which allows the removal of both organic and inorganic contaminants should be part of a standard operating procedure during the validation process. In this manner, pursuing false positives as genuine hits results in most cases in flat and irrational SAR, and can be avoided.

Finally, our survey of the PL^Pro^ screening literature highlighted some interesting trends. Most reported screening campaigns do have an additive included in the buffer; however, this is most often a single additive, DDT, which doesn’t fully prevent the inhibitory effects of copper. In general, we conclude that only 2 of the 28 studies reported account for potential metal contamination of both copper and palladium. Further, it has been many years since the original PAINS publication[55] but several reported nsp3 hits appear likely to be driving false-positives through fluorescence (dye-like chemical structures) and other PAINS mechanisms. Subsequently there is a greater need for adequate counter-screens in the nsp3 assay field and also increased awareness of false positive mechanisms more broadly.

Whilst this is what we have observed for screening of nsp3, we have found contamination with metal ions having effects in at least one other HTS screen carried out in the DDU on a completely different target. This suggests that this issue may be more general and the risk of the effects of metal ions commonly used in organic chemical synthesis (such as Cu and Pd) checked during assay development.

## Data availability

Synthetic procedures, chemical characterisation and further biochemical assay data are reported in the Supporting Information.

## Acknowledgements

This work was kindly funded by the Gates Foundation [INV-016131]. The authors kindly thank the DDU Compound Management Team (Alex Cookson, Kirsty Cookson, Fraser Hughes, Steve Bell) and the MRC PPU.

