## Supplemental Information for "Experience from an HTS campaign: Investigation of susceptibility and rescue of SARS-CoV-2 nsp3 protease assay from metal contamination"

### SUPPORTING INFORMATION

### 1. Chemical Synthesis

#### 1.1. Solvents and Reagents

All starting materials and solvents were obtained from standard chemical suppliers. Dry solvents were purchased in sure sealed bottles stored over molecular sieves. Organic solutions were routinely dried over anhydrous sodium sulfate or magnesium sulfate.

#### 1.2. Chromatography

Thin-layer chromatography was performed on pre-coated silica gel plates (Kieselgel 60 F254, BDH). Visualisation of the developed chromatogram was achieved by using fluorescence quenching and/or by staining with potassium permanganate. Column chromatography purification was performed on prepacked silica gel (230–400 mesh, 40–63 µm) cartridges.

#### 1.3. NMR Spectroscopy

All $^{1}H$ NMR spectra were recorded on a Brucker Avance III 400 MHz. Chemical shifts (δ) are expressed in ppm recorded using the residual undeuterated solvent as the internal reference in all cases. Signal splitting patterns are described as singlet (s), doublet (d), triplet (t), quartet (q), multiplet (m), broad (br), or a combination. Coupling constants (J) are quoted to the nearest 0.1 Hz.

#### 1.4. Reverse Phase HPLC conditions for the LCMS Analytical Methods

**Method 1:** SHIMADZU LCMS-2020 Kinetex EVO C18 2.1 × 30 mm, 5 µm at 50 ℃; Mobile Phase: A: 0.0375% TFA in water (v/v); B: 0.01875% TFA in Acetonitrile; flow rate held at 1.5 mL/min; eluted with the mobile phase over 1.55 min employing UV detection at 220&254 nm. Gradient information: 0-0.80 min, ramped from 95% A-5% B to 5% A-95% B; 0.80-1.20 min, held at 5% A-95% B; 1.20-1.21 min, returned to 95% A-5% B, 1.21-1.55 min, held at 95% A-5% B.
**Method 2:** SHIMADZU LCMS-2020 Kinetex EVO C18 2.1 × 30 mm, 5 µm at 50 ℃; Mobile Phase: A: 0.0375% TFA in water (v/v); B: 0.01875% TFA in Acetonitrile; flow rate held at 1.5 mL/min; eluted with the mobile phase over 1.55 min employing UV detection at 220&254 nm. Gradient information: 0-0.80 min, ramped from 100% A-0% B to 40% A-60% B; 0.80-1.20 min, held at 40% A-60% B; 1.20-1.21 min, returned to 100% A-0% B, 1.21-1.55 min, held at 100% A-0% B.
**Method 3:** Agilent 1200\G6110A Kinetex EVO C18 2.1 × 30 mm, 5 µm at 40 ℃; Mobile Phase: A: 0.0375% TFA in water (v/v); B: 0.01875% TFA in Acetonitrile; flow rate held at 1.5 mL/min; eluted with the mobile phase over 1.50 min employing UV detection at 220&254 nm. Gradient information: 0.01-0.80 min, ramped from 95% A-5% B to 5% A-95% B; 0.80-1.20 min, held at 5% A-95% B; 1.20-1.21 min, returned to 95% A-5% B, 1.21-1.5 min, held at 95% A-5% B.

##### 1.5. Reverse Phase HPLC conditions for the HPLC Analytical Methods

**<u>Method 1</u>:** SHIMADZU LC-20AB XBridge C18 2.1×50 mm, 5 µm at 40 ℃; Mobile Phase: A: 0.025% $NH_3 \cdot H_2O$ in water (v/v); B: Acetonitrile; flow rate held at 0.8 mL/min; eluted with the mobile phase over 4.00 min employing UV detection at 220&254 nm. Gradient information: 0-2.40 min, ramped from 90% A-10% B to 20% A-80% B; 2.40-3.20 min, held at 20% A-80% B; 3.20-3.20 min, returned to 90% A-10% B, 3.20-4.00 min, held at 90% A-10% B.
**<u>Method 2</u>:** SHIMADZU LC-20AB Kinetex EVO C18 4.6×50 mm, 5 µm at 50 ℃; Mobile Phase: A: 0.0375% TFA in water (v/v); B: 0.01875% TFA in Acetonitrile; flow rate held at 1.5 mL/min; eluted with the mobile phase over 4.00 min employing UV detection at 220&254 nm. Gradient information: 0-2.40 min, ramped from 90% A-10% B to 20% A-80% B; 2.40-3.70 min, held at 20% A-80% B; 3.70-3.71 min, returned to 90% A-10% B, 3.71-4.00 min, held at 90% A-10% B.

##### 1.6. Reverse Phase HPLC conditions for the HRMS Analytical Methods

**<u>Method 1</u>:** Agilent 1290 LC & Agilent G6530B Q-TOF Agilent ZORBAX Extend-C18 2.1 × 50 mm, 5 µm at 40 ℃; Mobile Phase: A: 0.0375% TFA in water (v/v); B: 0.0188% TFA in Acetonitrile; flow rate held at 0.8 mL/min; eluted with the mobile phase over 4.50 min employing DAD detection. Gradient information: 0-0.40 min, held at 0% A-100% B; 0.40-3.40 min, ramped from 0% A-100% B to 10% A-90% B; 3.40-3.90 min, ramped from 10% A-90% B to 0% A-100% B; 3.91-4.00 min, returned to 0% A-100% B, 4.00-4.50 min, held at 0% A-100% B (flow rate held at 1.0 mL/min).

##### 1.7. Preparative Reverse Phase HPLC conditions

**<u>Method 1</u>:** column: Waters Xbridge 150 × 25 mm × 5 µm; mobile phase: [water ($NH_4HCO_3$)-ACN]; B%: 2%-32%, 10 min.

##### 1.8. Synthetic procedures

###### 1.8.1 Synthesis of compound 2

**Scheme S1.** Synthetic route towards compound 2

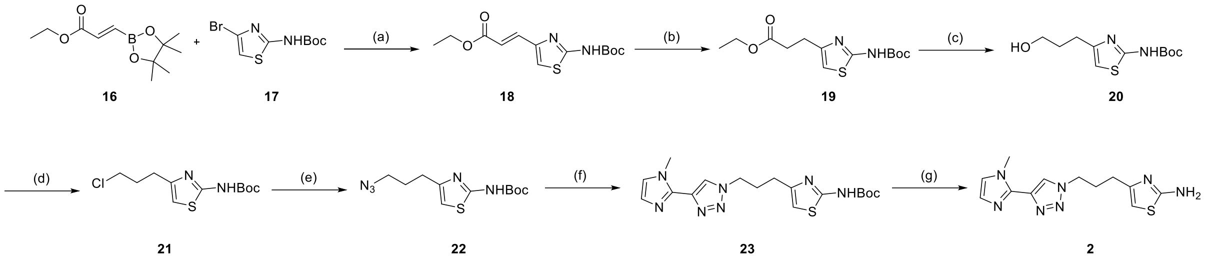

(a) $Na_2CO_3$, Pd(dppf)$Cl_2$, 1,4-dioxane, 80 °C, 16 h, 48%; (b) $H_2$, Pd/C, MeOH, 15 °C, 16 h, 67%; (c) $LiAlH_4$, THF, 0 °C, 1 h, 99%; (d) $SOCl_2$, DCM, 0 - 15 °C, 16h, 52%;(e) $TMSN_3$, TBAF, MeCN, 80 °C; (f) 2-ethynyl-1-methyl-imidazole, $CuSO_4{\cdot}5H_2O$, t-BuOH/$H_2O$, , 15 °C, 5 h, 48%; (g) HCl, MeOH, RT, 2 h, 88%.

#### Ethyl (E)-3-[2-(tert-butoxycarbonylamino)thiazol-4-yl]prop-2-enoate (18)

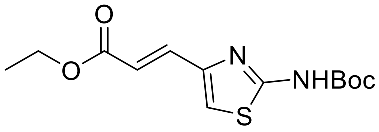

To a mixture of ethyl (E)-3-(4,4,5,5-tetramethyl-1,3,2-dioxaborolan-2-yl)prop-2-enoate **16** (506 mg, 2.24 mmol) and *tert*-butyl *N*-(4-bromothiazol-2-yl)carbamate **17** (500 mg, 1.79 mmol) in 1,4-dioxane (9 mL) and $H_2O$ (1 mL) was added $Na_2CO_3$ (380 mg, 3.58 mmol) and Pd(dppf)$Cl_2$ (131 mg, 0.18 mmol). The reaction mixture was stirred for 16 h at 80 °C under an inert atmosphere and then concentrated under vacuum. The residue was purified by flash column chromatography (petroleum ether/ethyl acetate, gradient 0-17%) to afford compound **18** (260 mg, 48% yield) as a light-yellow oil. LC MS ($ES^+$, method 1) *m/z* 243.1 $[M\text{-}tBu+H]^+$.

#### Ethyl 3-[2-(tert-butoxycarbonylamino)thiazol-4-yl]propanoate (19)

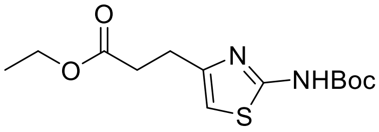

To a solution of compound **19** (250 mg, 838 µmol) in MeOH (7.5 mL) was added Pd/C (200 mg, 167 µmol, 10% wt), and the mixture was stirred for 16 h at 15 °C under a $H_2$ balloon (15 psi). The reaction mixture was filtered, and the filtrate was concentrated under vacuum to afford compound **19** (170 mg, 67% yield) as colourless oil. The isolated product was used in the following step without any further purification. LC MS ($ES^+$, method 3) *m/z* 301.1 $[M+H]^+$.

#### *tert*-Butyl N-[4-(3-hydroxypropyl)thiazol-2-yl]carbamate (20)

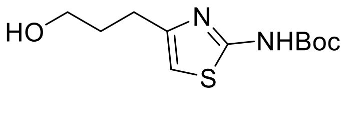

To a solution of compound **19** (210 mg, 699 µmol) in THF (5 mL) at 0 °C under an inert atmosphere, was added dropwise $LiAlH_4$ (839 µL , 1 M in THF), and left to stir at 0 °C for 1 h. The reaction mixture was gradually quenched with $H_2O$ (7.58 µL), 15% aq. NaOH (7.58 µL) and $H_2O$ (22.5 µL) successively. The mixture was stirred for 30 min, filtered, and rinsed with THF (3 x 5 mL). The filtrate was concentrated under vacuum to afford compound **20** (180 mg, 99% yield) as a light-yellow oil. The isolated product was used in the following step without any further purification. LC MS ($ES^+$, method 3) *m/z* 259.1 $[M+H]^+$.

**tert-Butyl N-[4-(3-chloropropyl)thiazol-2-yl]carbamate (21)**

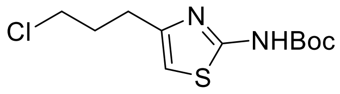

To a solution of compound **20** (180 mg, 0.70 mmol) in DCM (7.5 mL) at 0 °C was added $SOCl_2$ (253 µL, 3.48 mmol). The reaction mixture was stirred for 16 h at 15 °C and then concentrated under vacuum. The residue was purified by reversed phase HPLC (0.1% formic acid in $H_2O$) to afford compound **21** (100 mg, 52% yield) as a brown oil. LC MS ($ES^+$, method 3) *m/z* 221.1 $[M-C_4H_8+H]^+$.

***tert*-Butyl N-[4-(3-azidopropyl)thiazol-2-yl]carbamate (22)**

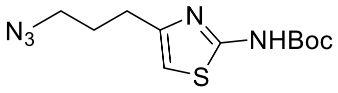

To a solution of compound **21** (100 mg, 361 µmol) and TBAF (722 µL, 1 M in THF) in $CH_3CN$ (2.5 mL) was added $TMSN_3$ (95.0 µL, 722 µmol) at 15 °C. The reaction mixture was stirred for 16 h at 80 °C and then concentrated under vacuum. The residue was diluted with water (20 mL) and the mixture was extracted with EtOAc (2 x 30 mL) and then washed with brine (2 x 30 mL). The organic layer was dried, filtered, and the filtrate concentrated under vacuum. The residue was purified by reversed phase HPLC (0.1% formic acid in $H_2O$/MeCN, gradient 0-100%) to afford compound **22** (40.0 mg, 39% yield) as a light-yellow oil. LC MS ($ES^+$, method 1) *m/z* 228.1 $[M+H-C_4H_8]^+$.

**tert-Butyl N-[4-[3-[4-(1-methylimidazol-2-yl)triazol-1-yl]propyl]thiazol-2-yl]carbamate (23)**

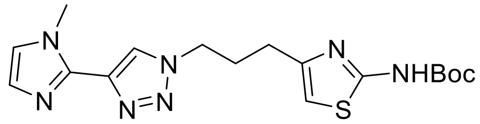

To a suspension of compound **22** (30.0 mg, 106 µmol) in $H_2O$ (1 mL) and t-BuOH (1 mL) was added 2-ethynyl-1-methyl-imidazole (11.2 mg, 106 µmol), followed by sodium ascorbate (10.9 µL, 1.06

µmol) and $CuSO_4 \cdot 5H_2O$ (264 mg, 1.06 µmol). The reaction mixture was stirred for 5 h at 15 °C and then concentrated under vacuum. The crude product was purified by reversed phase HPLC (0.1% formic acid in $H_2O$) to afford compound **23** (20.0 m, 48% yield) as a light-yellow solid. LC MS ($ES^+$, method 2) *m/z* 390.1 $[M+H]^+$.

**4-[3-[4-(1-Methylimidazol-2-yl)triazol-1-yl]propyl]thiazol-2-amine (2)**

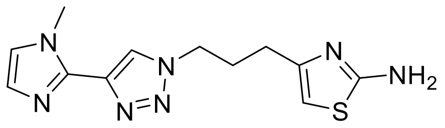

To a solution of compound **23** (35.0 mg, 89.8 µmol) in HCl (898 µL, 4 M in MeOH) was added DCM (1 mL). The reaction mixture was stirred at RT for 2 h. The residual aqueous solution was lyophilized in $H_2O$ to afford compound **2** (26.9 mg, 88% yield) as a brown solid. HPLC (method 1) 95.7% purity; HRMS (method 1) *m/z* $[M+H]^+$ calcd for $C_{12}H_{15}N_7S$ 290.1188; found 290.1184; $^1H$ NMR (400 MHz, $D_2O$): $\delta$ 8.69 (s, 1H), 7.53 (s, 2H), 6.40 (s, 1H), 4.65 – 6.62 (m, 2H), 4.01 (s, 3H), 2.71 - 2.64 (m, 2H), 2.38 - 2.31 (m, 2H).

#### 2. Biochemical Assay Materials and Methods

##### 2.1. Materials

Enzyme: SARS-CoV-2 nsp3 (179-1329) Twinstrep, MRC-PPU, University of Dundee, DU67831; Substrate: Z-RLRGG-AMC, Bachem, 4027158; Assay Buffer: HEPES, Formedium HEPES10; NaCl, Sigma S7653; BSA, Sigma A7906; NP-40*, Abcam ab142227; Plates: Black 384-well, low-volume assay plates, Greiner 784900; Additives: (1) metal compounds CuI, $Cu(OAc)_2$, $Pd(PPh_3)_4$, $Pd(OAc)_2$; (2) chelators EDTA; or EGTA; or GSH; or DTT; or TCEP.

##### 2.2. High Throughput Screen

25nM SARS-CoV-2 nsp3 (final assay concentration) in 5 µl assay buffer (40 mM HEPES, 100 mM NaCl, 0.005% BSA, 0.01% NP-40, pH7.5) was added to compound-stamped 384 well assay plates. Compounds were stamped in 100 nl DMSO giving a final [DMSO] of 1%. The assay was started with the addition of 30 µM (final assay concentration) Z-RLRGG-AMC substrate (5 µl) then the plates were covered and incubated at room temp (20$^{o}$C) for 3 hours. Plates were then read on a BMG Pherastar using a 350/450 fluorescence filter. Data was analysed in Activity Base.

##### 2.3. SARS-CoV-2-nsp3 enzymatic assay in the presence of metal compounds

25nM SARS-CoV-2 nsp3 (final assay concentration) in 5 µl assay buffer (40 mM HEPES, 100 mM NaCl, 0.005% BSA, 0.01% NP-40, pH7.5) was added to metal compound-stamped 384 well assay plates. Metal compounds were stamped in 100 nl DMSO giving a final [DMSO] of 1%. All metal compounds were tested at 0.5 nM-10 µM final assay concentration and all samples were prepared fresh on the day of assay. The assay was started with the addition of 30 µM (final assay concentration) Z-RLRGG-AMC substrate (5 µl) then the plates were covered and incubated at room temp (20$^{o}$C) for 3 hours. Plates were then read on a BMG Pherastar using a 350/450 fluorescence filter. Data was analysed using XLFit and GraphFit.

##### 2.4. SARS-CoV-2-nsp3 enzymatic assay in the presence of metal compounds and chelators/reducers

25nM SARS-CoV-2 nsp3 (final assay concentration) in 5 µl assay buffer (40 mM HEPES, 100 mM NaCl, 0.005% BSA, 0.01% NP-40, pH7.5) containing 0 mM, 0.5 mM, 1 mM or 5 mM EDTA/EGTA/GSH/DTT/TCEP was added to metal compound-stamped 384 well assay plates. Metal compounds were stamped in 100 nl DMSO giving a final [DMSO] of 1%. All metal compounds were tested at 0.5 nM-10 µM final assay concentration and all samples were prepared fresh on the day of assay. The assay was started with the addition of 30 µM (final assay concentration) Z-RLRGG-AMC substrate (5 µl) then the plates were covered and incubated at room temp (20$^{o}$C) for 3 hours. Plates were then read on a BMG Pherastar using a 350/450 fluorescence filter. Data was analysed using XLFit and GraphFit.

##### 2.5. SARS-CoV-2-nsp3 enzymatic assay in the presence of chelators/reducers

25nM SARS-CoV-2 nsp3 (final assay concentration) in 5 µl assay buffer (40 mM HEPES, 100 mM NaCl, 0.005% BSA, 0.01% NP-40, pH7.5) containing 0 mM or 5 mM EDTA/DTT was added to compound-stamped 384 well assay plates. Compounds were stamped in 100 nl DMSO giving a final [DMSO] of 1%. The assay was started with the addition of 30 µM (final assay concentration) Z-RLRGG-AMC substrate (5 µl) then the plates were covered and incubated at room temp (20°C) for 3 hours. Plates were then read on a BMG Pherastar using a 350/450 fluorescence filter. Data was analysed in Activity Base.

#### 3. Additional Information

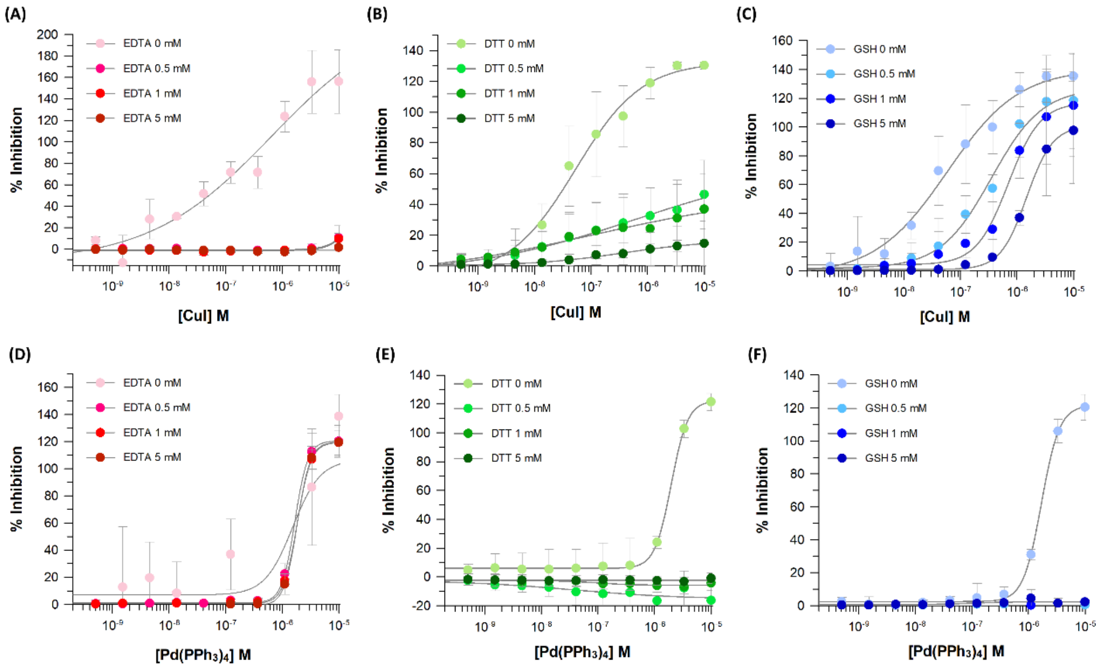

**Figure S1.** Inhibitory effect of CuI against SARS-CoV-2 nsp3 in the presence of (A) 0.5, 1 and 5 mM of EDTA, (B) 0.5, 1 and 5 mM of DTT, (C) 0.5, 1 and 5 mM of GSH. Inhibitory effect of [Pd(PPh$_3$)$_4$]. against SARS-CoV-2 nsp3 in the presence of (D) 0.5, 1 and 5 mM of EDTA, (E) 0.5, 1 and 5 mM of DTT, (F) 0.5, 1 and 5 mM of GSH. Data shown represent mean ± SDTV from three independent experiments.

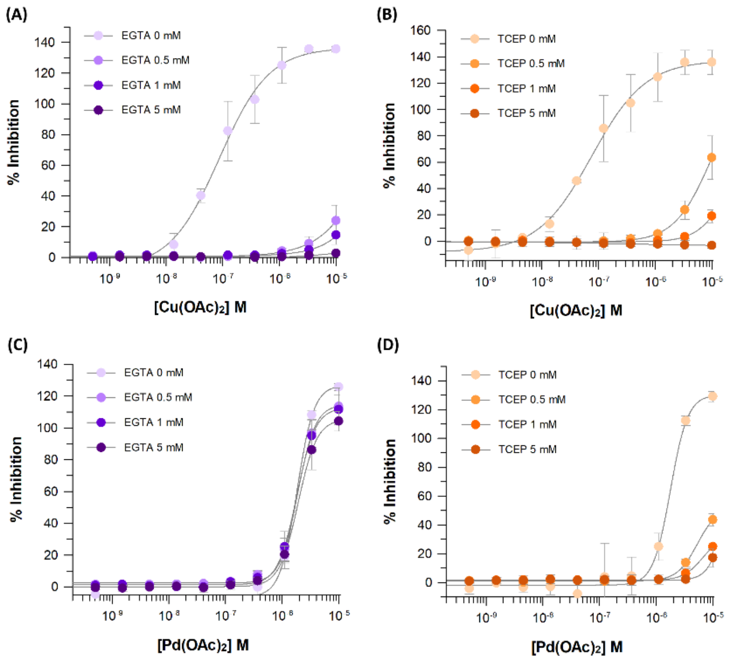

**Figure S2.** Inhibitory effect of $Cu(OAc)_2$ against SARS-CoV-2 nsp3 in the presence of (A) 0.5, 1 and 5 mM of EGTA, (B) 0.5, 1 and 5 mM of TCEP. Inhibitory effect of $Pd(OAc)_2$. against SARS-CoV-2 nsp3 in the presence of (C) 0.5, 1 and 5 mM of EGTA, (D) 0.5, 1 and 5 mM of TCEP. Data shown represent mean ± SDTV from three independent experiments.

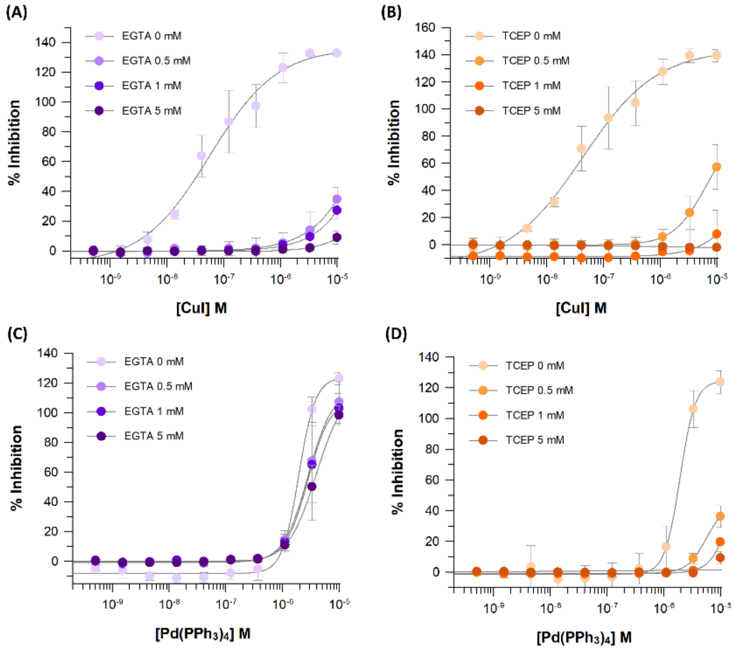

**Figure S3.** Inhibitory effect of CuI against SARS-CoV-2 nsp3 in the presence of (A) 0.5, 1 and 5 mM of EGTA, (B) 0.5, 1 and 5 mM of TCEP. Inhibitory effect of $[Pd(PPh_3)_4]$. against SARS-CoV-2 nsp3 in the presence of (C) 0.5, 1 and 5 mM of EGTA, (D) 0.5, 1 and 5 mM of TCEP. Data shown represent mean ± SDTV from three independent experiments.

### 4. Appendix

#### 4.1 QC Spectra for Compounds

**LCMS** Ethyl (E)-3-[2-(tert-butoxycarbonylamino)thiazol-4-yl]prop-2-enoate (**18**)

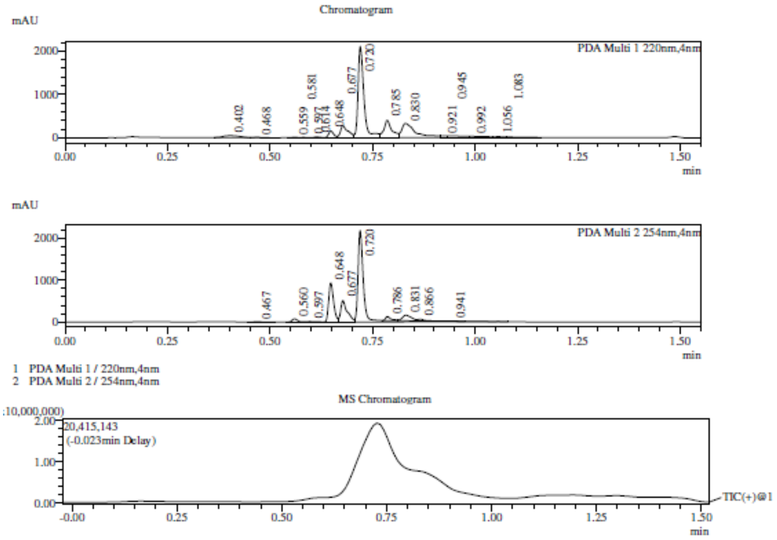

Integration Result

Peak Table

PDA Ch1 220nm

| Peak# | Ret. Time | Height | Height% | USP Width | Area | Area% |
|---|---|---|---|---|---|---|
| 1 | 0.402 | 47591 | 1.352 | 0.075 | 132380 | 3.051 |
| 2 | 0.468 | 8273 | 0.235 | 0.039 | 11397 | 0.263 |
| 3 | 0.559 | 12846 | 0.365 | 0.026 | 11198 | 0.258 |
| 4 | 0.581 | 9370 | 0.266 | 0.028 | 6955 | 0.160 |
| 5 | 0.597 | 7421 | 0.211 | 0.083 | 5950 | 0.137 |
| 6 | 0.614 | 19958 | 0.567 | 0.029 | 18969 | 0.437 |
| 7 | 0.648 | 156929 | 4.459 | 0.026 | 137079 | 3.159 |
| 8 | 0.677 | 278272 | 7.907 | 0.034 | 362707 | 8.359 |
| 9 | 0.720 | 2104252 | 59.795 | 0.026 | 2055682 | 47.377 |
| 10 | 0.785 | 403117 | 11.455 | 0.033 | 534549 | 12.320 |
| 11 | 0.830 | 329097 | 9.352 | 0.052 | 773747 | 17.832 |
| 12 | 0.921 | 42825 | 1.217 | 0.328 | 46243 | 1.066 |
| 13 | 0.945 | 39059 | 1.110 | 0.545 | 110201 | 2.540 |
| 14 | 0.992 | 28158 | 0.800 | 0.000 | 77648 | 1.790 |
| 15 | 1.056 | 20088 | 0.571 | 0.085 | 31325 | 0.722 |
| 16 | 1.083 | 11847 | 0.337 | 0.443 | 22986 | 0.530 |

Peak Table

PDA Ch2 254nm

| Peak# | Ret. Time | Height | Height% | USP Width | Area | Area% |
|---|---|---|---|---|---|---|
| 1 | 0.467 | 5290 | 0.133 | 0.042 | 8046 | 0.203 |
| 2 | 0.560 | 74653 | 1.875 | 0.026 | 68032 | 1.720 |
| 3 | 0.597 | 11248 | 0.283 | 0.040 | 16663 | 0.421 |
| 4 | 0.648 | 929433 | 23.345 | 0.024 | 777814 | 19.664 |
| 5 | 0.677 | 510423 | 12.821 | 0.030 | 572505 | 14.474 |
| 6 | 0.720 | 2175065 | 54.633 | 0.024 | 2161100 | 54.636 |
| 7 | 0.786 | 94388 | 2.371 | 0.025 | 91638 | 2.317 |
| 8 | 0.831 | 132692 | 3.333 | 0.047 | 206569 | 5.222 |
| 9 | 0.866 | 43811 | 1.100 | 0.035 | 47126 | 1.191 |
| 10 | 0.941 | 4261 | 0.107 | 0.047 | 5969 | 0.151 |

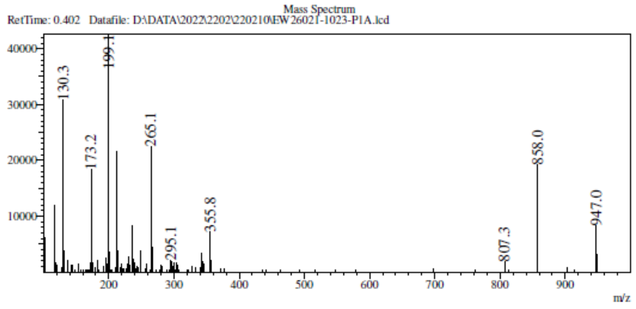

RetTime: 0.652   Datafile: D:\DATA\2022\2202\220210\EW26021-1023-P1A.lcd

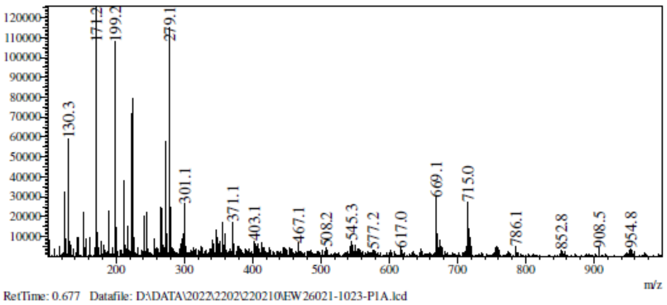

RetTime: 0.677   Datafile: D:\DATA\2022\2202\220210\EW26021-1023-P1A.lcd

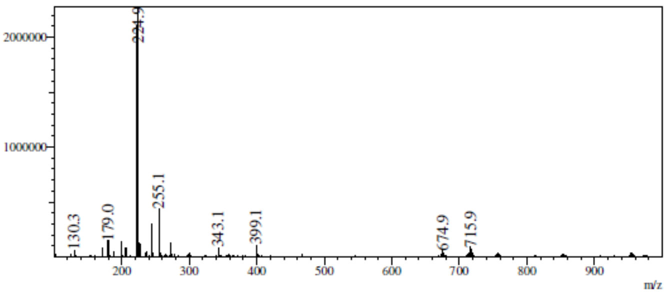

RetTime: 0.718   Datafile: D:\DATA\2022\2202\220210\EW26021-1023-P1A.lcd

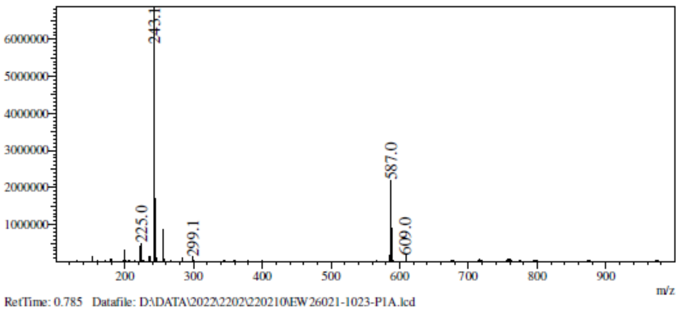

RetTime: 0.785   Datafile: D:\DATA\2022\2202\220210\EW26021-1023-P1A.lcd

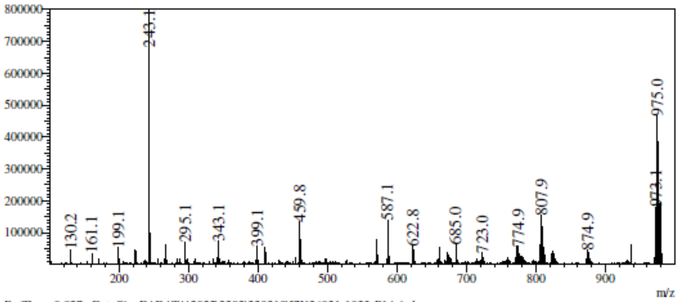

RetTime: 0.827   Datafile: D:\DATA\2022\2202\220210\EW26021-1023-P1A.lcd

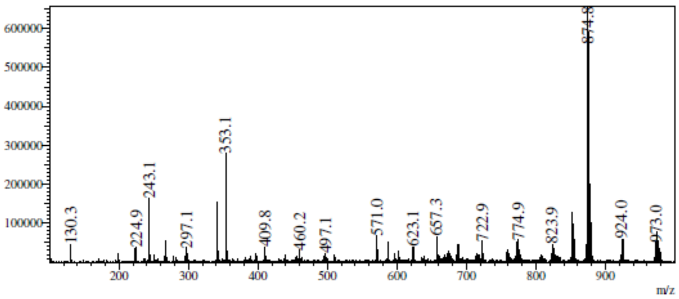

**LCMS** Ethyl 3-[2-(tert-butoxycarbonylamino)thiazol-4-yl]propanoate (**19**)

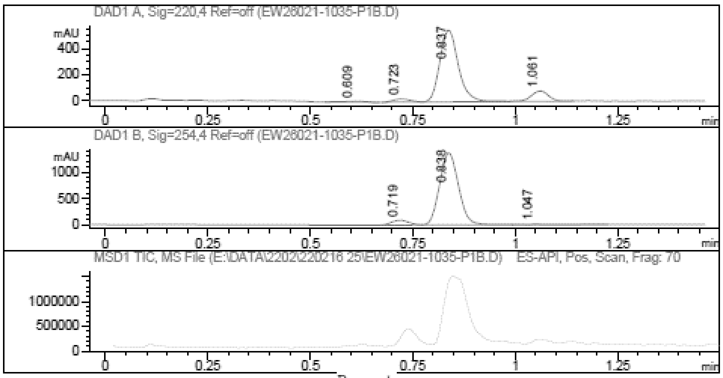

Report

Signal 1 : DAD1 A, Sig=220,4 Ref=off

| # | Meas. Ret. | Height | Width | Area | Area % |
|---|---|---|---|---|---|
| 1 | 0.609 | 7.011 | 0.055 | 27.097 | 1.295 |
| 2 | 0.723 | 24.102 | 0.046 | 70.218 | 3.356 |
| 3 | 0.837 | 552.203 | 0.051 | 1757.678 | 84.003 |
| 4 | 1.061 | 79.639 | 0.047 | 237.411 | 11.346 |

Signal 2 : DAD1 B, Sig=254,4 Ref=off

| # | Meas. Ret. | Height | Width | Area | Area % |
|---|---|---|---|---|---|
| 1 | 0.719 | 92.741 | 0.046 | 267.264 | 5.629 |
| 2 | 0.838 | 1369.569 | 0.052 | 4427.279 | 93.245 |
| 3 | 1.047 | 10.436 | 0.074 | 53.439 | 1.126 |

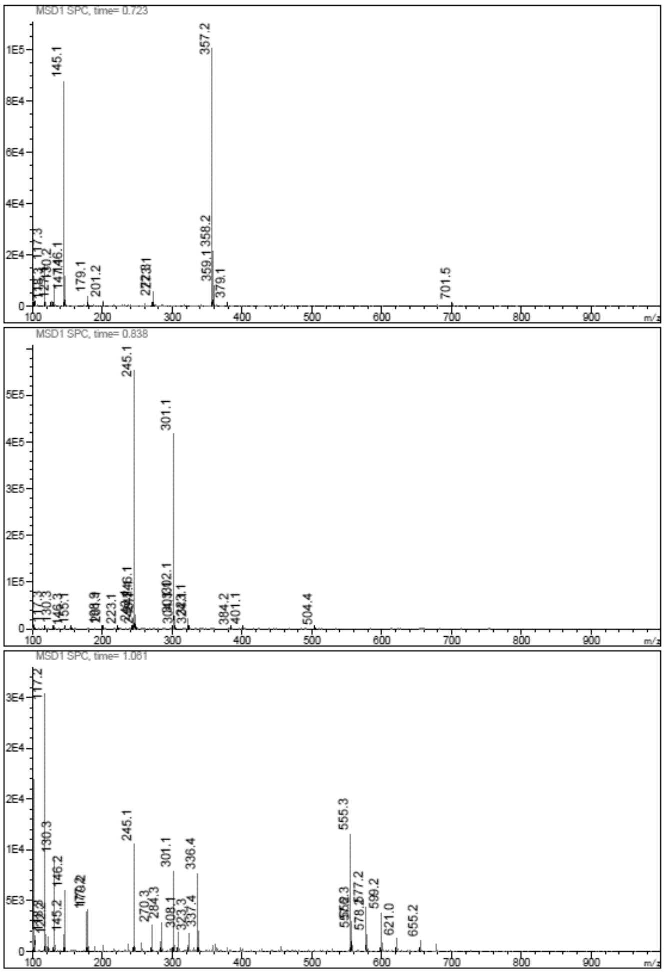

**LCMS** *tert*-Butyl N-[4-(3-hydroxypropyl)thiazol-2-yl]carbamate (**20**)

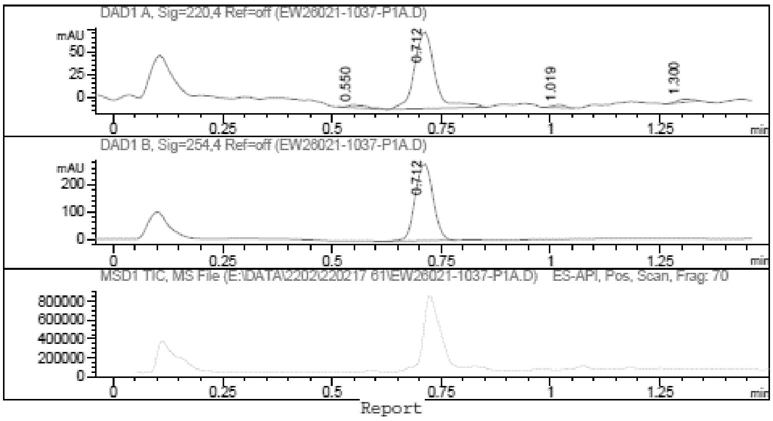

Report

Signal 1 : DAD1 A, Sig=220,4 Ref=off

| # | Meas. Ret. | Height | Width | Area | Area % |
|---|---|---|---|---|---|
| 1 | 0.550 | 3.242 | 0.042 | 10.064 | 3.317 |
| 2 | 0.712 | 84.796 | 0.050 | 277.955 | 91.616 |
| 3 | 1.019 | 3.346 | 0.036 | 6.791 | 2.238 |
| 4 | 1.300 | 3.503 | 0.041 | 8.582 | 2.829 |

Signal 2 : DAD1 B, Sig=254,4 Ref=off

| # | Meas. Ret. | Height | Width | Area | Area % |
|---|---|---|---|---|---|
| 1 | 0.712 | 283.974 | 0.045 | 806.756 | 100.000 |

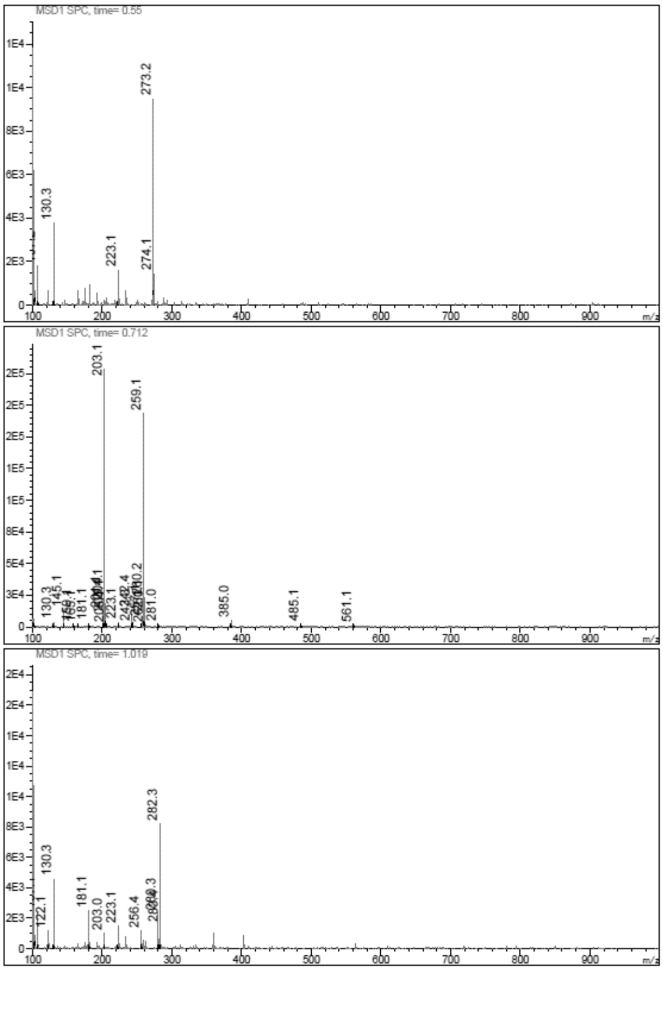

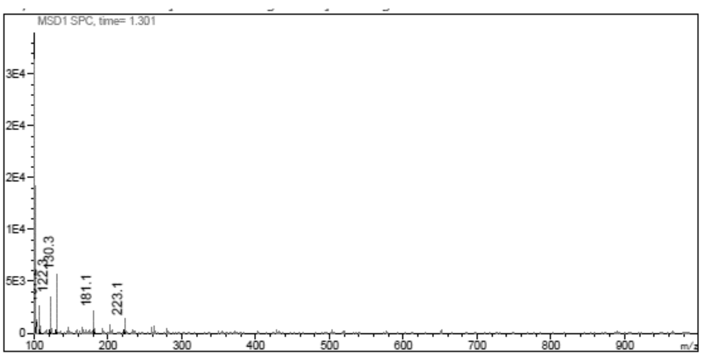

**LCMS** tert-Butyl N-[4-(3-chloropropyl)thiazol-2-yl]carbamate (**21**)

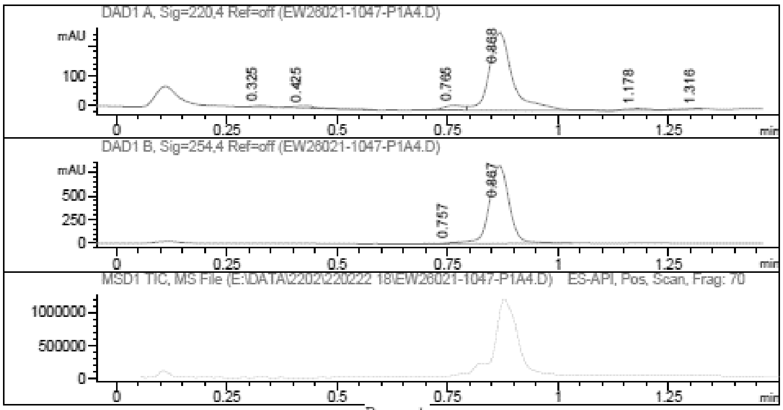

Report

Signal 1 : DAD1 A, Sig=220,4 Ref=off

| # | Meas. Ret. | Height | Width | Area | Area % |
|---|---|---|---|---|---|
| 1 | 0.325 | 4.899 | 0.033 | 11.328 | 1.039 |
| 2 | 0.425 | 7.573 | 0.060 | 30.941 | 2.839 |
| 3 | 0.765 | 16.238 | 0.048 | 50.241 | 4.610 |
| 4 | 0.868 | 262.807 | 0.055 | 967.428 | 88.771 |
| 5 | 1.178 | 6.178 | 0.057 | 22.734 | 2.086 |
| 6 | 1.316 | 2.502 | 0.039 | 7.133 | 0.654 |

Signal 2 : DAD1 B, Sig=254,4 Ref=off

| # | Meas. Ret. | Height | Width | Area | Area % |
|---|---|---|---|---|---|
| 1 | 0.757 | 13.616 | 0.038 | 35.322 | 1.343 |
| 2 | 0.867 | 830.924 | 0.048 | 2594.303 | 98.657 |

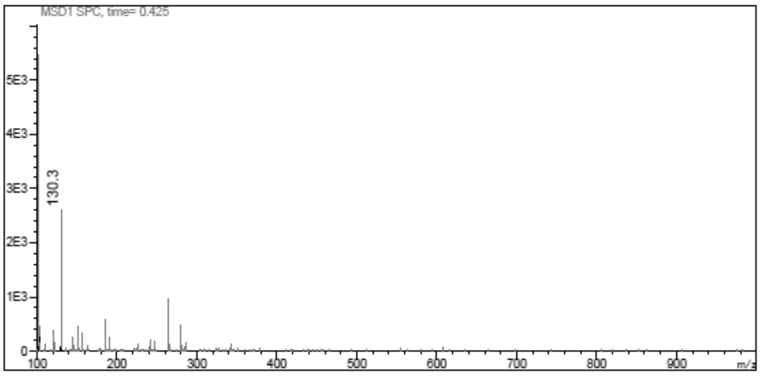

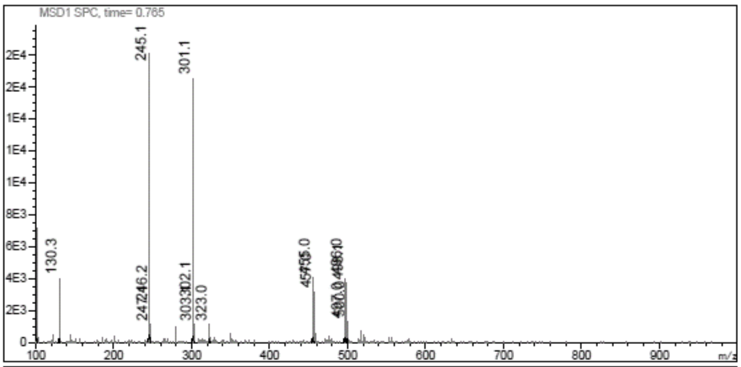
MSD1 SPC, time= 0.765
245.1
301.1
130.3
323.0
m/z

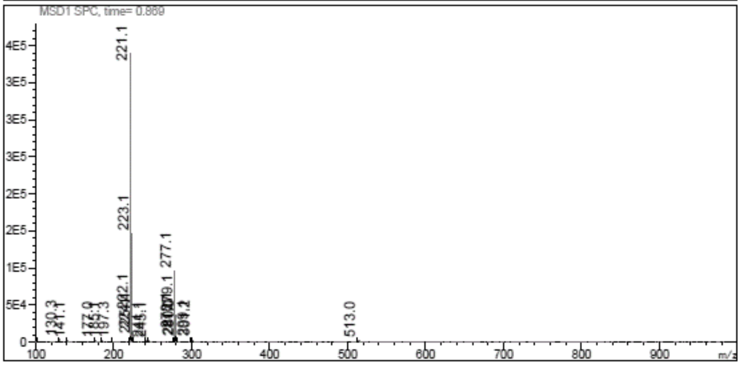
MSD1 SPC, time= 0.869
221.1
223.1
277.1
130.3
513.0
m/z

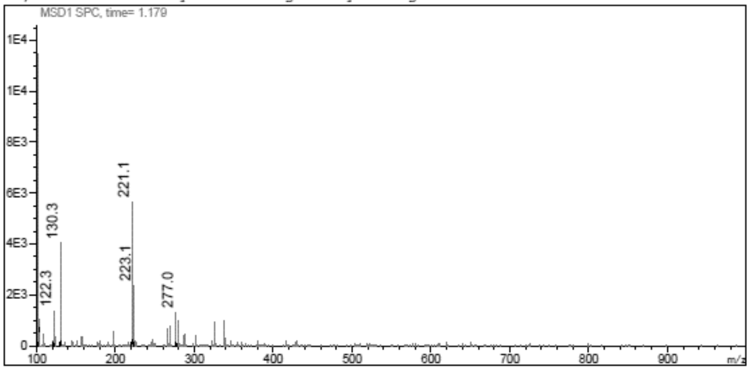
MSD1 SPC, time= 1.179
221.1
130.3
223.1
122.3
277.0
m/z

**LCMS** *tert*-Butyl N-[4-(3-azidopropyl)thiazol-2-yl]carbamate (**22**)

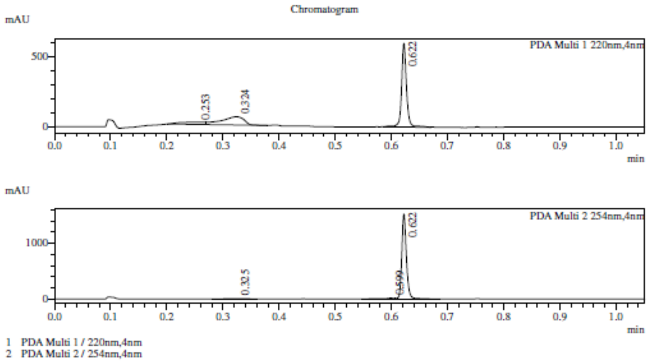

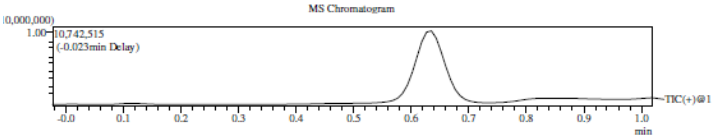

Integration Result

Peak Table

PDA Ch1 220nm

| Peak# | Ret. Time | Height | Height% | USP Width | Area | Area% |
|---|---|---|---|---|---|---|
| 1 | 0.253 | 16182 | 2.446 | 0.000 | 51544 | 8.852 |
| 2 | 0.324 | 58796 | 8.889 | 0.071 | 163514 | 28.079 |
| 3 | 0.622 | 586492 | 88.665 | 0.015 | 367266 | 63.069 |

Peak Table

PDA Ch2 254nm

| Peak# | Ret. Time | Height | Height% | USP Width | Area | Area% |
|---|---|---|---|---|---|---|
| 1 | 0.325 | 3017 | 0.201 | 0.057 | 7018 | 0.791 |
| 2 | 0.599 | 15003 | 0.999 | 0.024 | 16216 | 1.828 |
| 3 | 0.622 | 1483340 | 98.800 | 0.015 | 863726 | 97.380 |

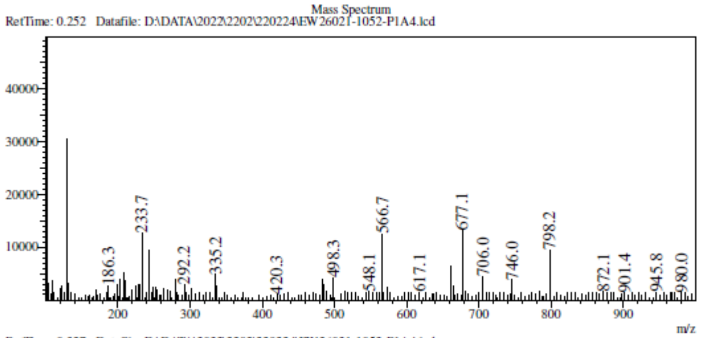

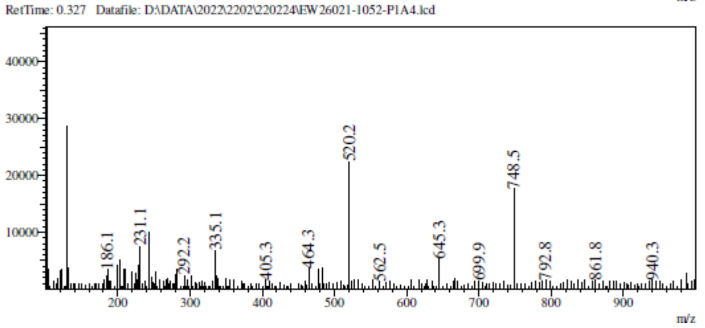

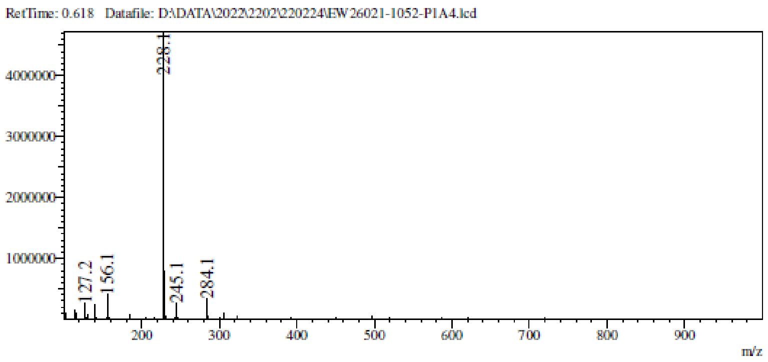

**LCMS** *tert*-Butyl N-[4-[3-[4-(1-methylimidazol-2-yl)triazol-1-yl]propyl]thiazol-2-yl]carbamate (**23**)

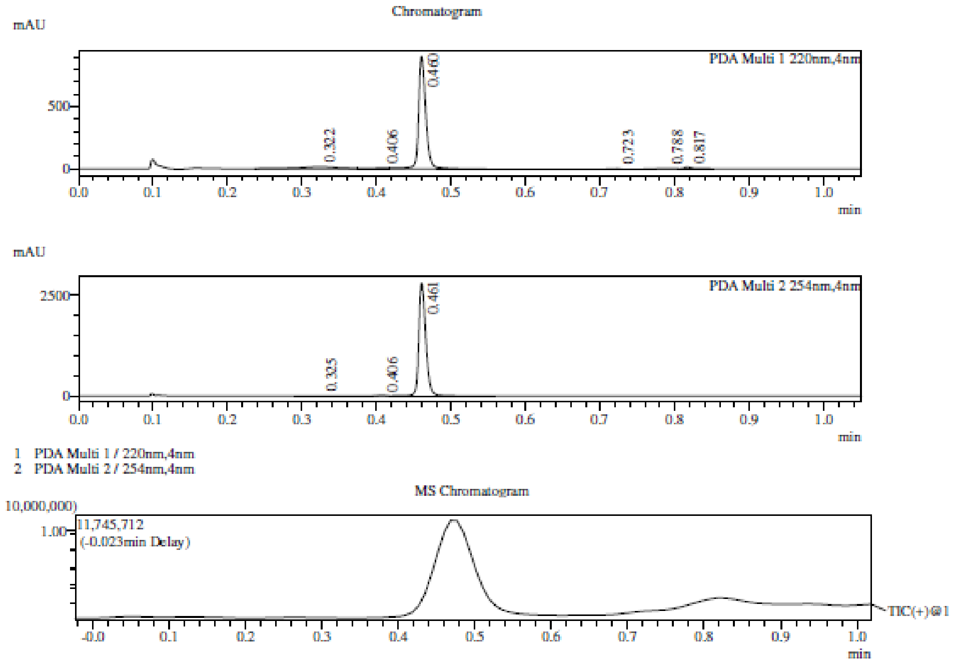

Integration Result

Peak Table

PDA Ch1 220nm

| Peak# | Ret. Time | Height | Height% | USP Width | Area | Area% |
|---|---|---|---|---|---|---|
| 1 | 0.322 | 16729 | 1.787 | 0.085 | 63392 | 8.688 |
| 2 | 0.406 | 9241 | 0.987 | 0.072 | 15655 | 2.146 |
| 3 | 0.460 | 887268 | 94.782 | 0.017 | 621762 | 85.218 |
| 4 | 0.723 | 5200 | 0.555 | 0.017 | 5999 | 0.822 |
| 5 | 0.788 | 2690 | 0.287 | 0.331 | 6841 | 0.938 |
| 6 | 0.817 | 14991 | 1.601 | 0.020 | 15966 | 2.188 |

Peak Table

PDA Ch2 254nm

| Peak# | Ret. Time | Height | Height% | USP Width | Area | Area% |
|---|---|---|---|---|---|---|
| 1 | 0.325 | 3199 | 0.115 | 0.077 | 12322 | 0.662 |
| 2 | 0.406 | 8560 | 0.309 | 0.023 | 7079 | 0.380 |
| 3 | 0.461 | 2759364 | 99.576 | 0.017 | 1841727 | 98.958 |

Mass Spectrum

RetTime: 0.318 Datafile: D:\DATA\2022\2203\220301\EW26021-1058-P1A1.lcd

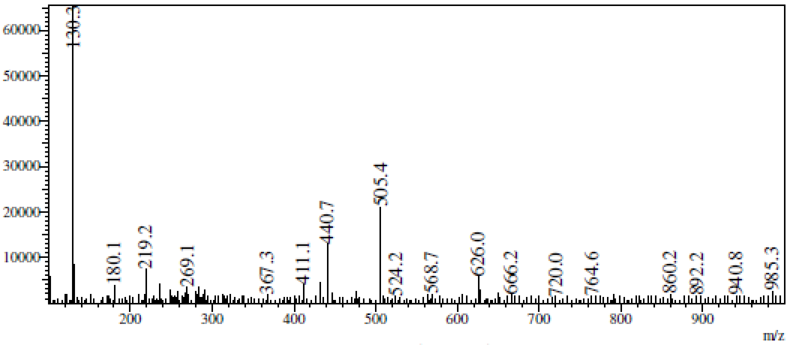

RetTime: 0.402 Datafile: D:\DATA\2022\2203\220301\EW26021-1058-P1A1.lcd

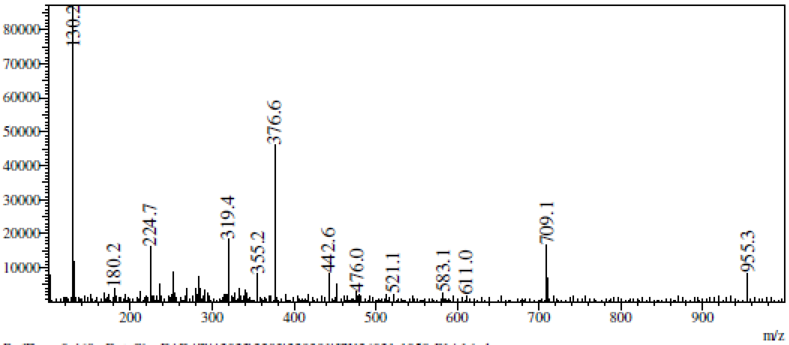

RetTime: 0.460 Datafile: D:\DATA\2022\2203\220301\EW26021-1058-P1A1.lcd

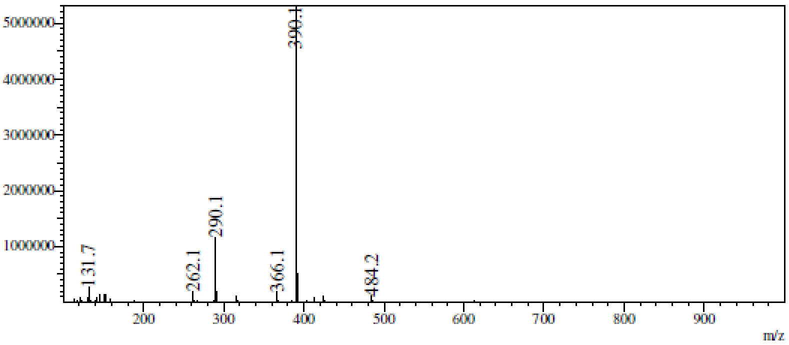

RetTime: 0.727 Datafile: D:\DATA\2022\2203\220301\EW26021-1058-P1A1.lcd

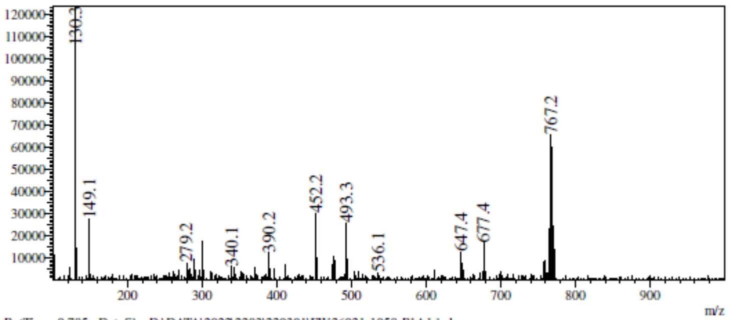

RetTime: 0.785 Datafile: D:\DATA\2022\2203\220301\EW26021-1058-P1A1.lcd

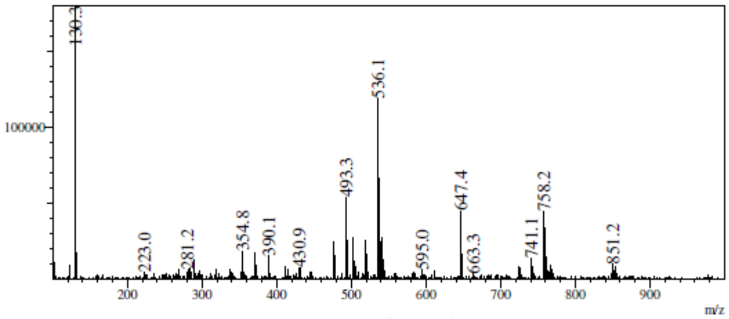

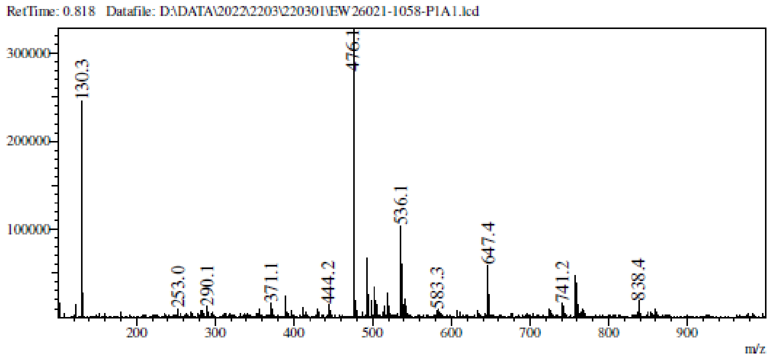

**$^1$H NMR** 4-[3-[4-(1-Methylimidazol-2-yl)triazol-1-yl]propyl]thiazol-2-amine (**2**)

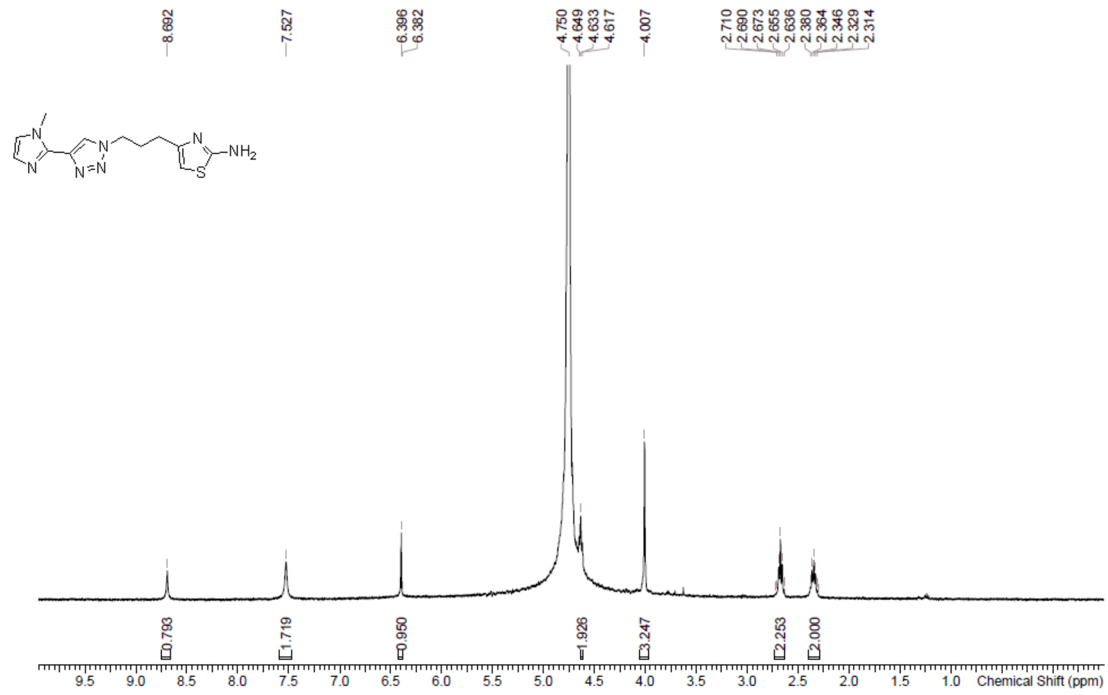

**HPLC** 4-[3-[4-(1-Methylimidazol-2-yl)triazol-1-yl]propyl]thiazol-2-amine (**2**)

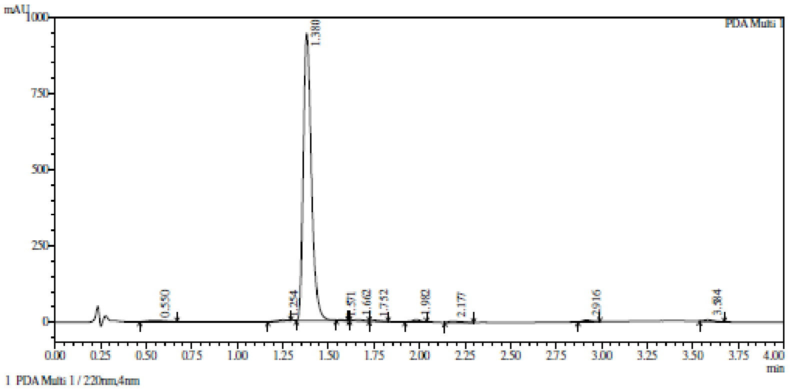

Integration result

PeakTable

PDA Ch1 220nm

| Peak# | Ret. Time | USP Width | Resolution | Height | Area | Area % |
|---|---|---|---|---|---|---|
| 1 | 0.550 | 0.173 | 0.000 | 3974 | 25933 | 0.836 |
| 2 | 1.254 | 0.091 | 5.338 | 2975 | 10659 | 0.344 |
| 3 | 1.380 | 0.081 | 1.472 | 941715 | 2969097 | 95.691 |
| 4 | 1.571 | 0.057 | 2.781 | 1386 | 2913 | 0.094 |
| 5 | 1.662 | 0.098 | 1.169 | 4101 | 13380 | 0.431 |
| 6 | 1.752 | 0.077 | 1.037 | 2948 | 8452 | 0.272 |
| 7 | 1.982 | 0.075 | 3.018 | 7379 | 21883 | 0.705 |
| 8 | 2.177 | 0.139 | 1.825 | 4104 | 18464 | 0.595 |
| 9 | 2.916 | 0.082 | 6.709 | 5375 | 16836 | 0.543 |
| 10 | 3.584 | 0.090 | 7.790 | 4532 | 15194 | 0.490 |
| Total | | | | 978490 | 3102813 | 100.000 |

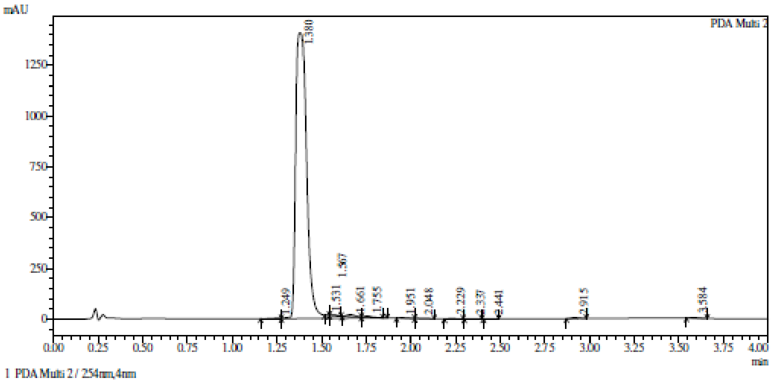

Integration result

PeakTable

PDA Ch2 254nm

| Peak# | Ret. Time | USP Width | Resolution | Height | Area | Area % |
|---|---|---|---|---|---|---|
| 1 | 1.249 | 0.213 | 0.000 | 2792 | 11164 | 0.184 |
| 2 | 1.380 | 0.093 | 0.853 | 1408042 | 5939074 | 97.895 |
| 3 | 1.531 | 0.000 | 0.000 | 1029 | 1295 | 0.021 |
| 4 | 1.567 | 0.062 | 0.000 | 4484 | 10774 | 0.178 |
| 5 | 1.661 | 0.095 | 1.189 | 10297 | 33852 | 0.558 |
| 6 | 1.755 | 0.076 | 1.100 | 6264 | 17974 | 0.296 |
| 7 | 1.951 | 0.087 | 2.405 | 2376 | 8132 | 0.134 |
| 8 | 2.048 | 0.106 | 1.013 | 1594 | 6544 | 0.108 |
| 9 | 2.229 | 0.090 | 1.848 | 1634 | 5526 | 0.091 |
| 10 | 2.337 | 0.083 | 1.259 | 1083 | 3338 | 0.055 |
| 11 | 2.441 | 0.068 | 1.383 | 1392 | 3590 | 0.059 |
| 12 | 2.915 | 0.082 | 6.327 | 4744 | 14537 | 0.240 |
| 13 | 3.584 | 0.086 | 7.966 | 3398 | 10981 | 0.181 |
| Total | | | | 1449129 | 6066782 | 100.000 |

**HRMS** 4-[3-[4-(1-Methylimidazol-2-yl)triazol-1-yl]propyl]thiazol-2-amine (**2**)

**Compound Table**

| Label | Tgt Score | Mass Error (ppm) | Tgt Formula | Obs. RT | Ref. Mass | Obs. Mass |
|---|---|---|---|---|---|---|
| Cpd 1: C12 H15 N7 S; 1.103 | 99.11 | 0.55 | C12 H15 N7 S | 1.103 | 289.111 | 289.1111 |

| Obs. *m/z* | Obs. RT | Obs. Mass | Tgt Formula | Tgt Mass | Tgt Mass Error (ppm) | Find Cpds Algorithm |
|---|---|---|---|---|---|---|
| 290.1184 | 1.103 | 289.1111 | C12 H15 N7 S | 289.111 | 0.55 | Find by Formula |

**Compound Chromatograms**

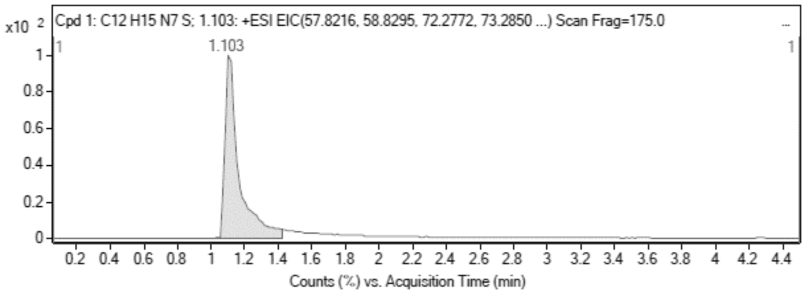

**MS Zoomed Spectrum**

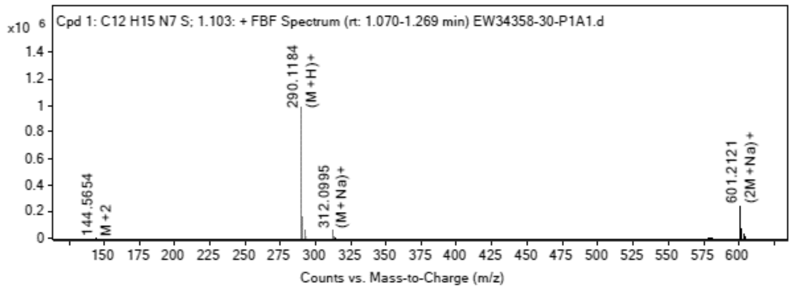

**MS Spectrum Peak List**

| Obs. *m/z* | Charge | Abund | Ion/Isotope |
|---|---|---|---|
| 290.1184 | 1 | 986351.63 | (M+H)+ |
| 291.12 | 1 | 159996.56 | (M+H)+ |
| 292.1153 | 1 | 52298.26 | (M+H)+ |
| 312.0995 | 1 | 54301.61 | (M+Na)+ |
| 313.1017 | 1 | 8462.2 | (M+Na)+ |
| 314.0968 | 1 | 2829.1 | (M+Na)+ |
| 601.2121 | 1 | 236698.11 | (2M+Na)+ |
| 602.2139 | 1 | 76042.87 | (2M+Na)+ |
| 603.2102 | 1 | 31429.46 | (2M+Na)+ |
| 604.2101 | 1 | 7531.87 | (2M+Na)+ |

**MS Zoomed Spectrum**

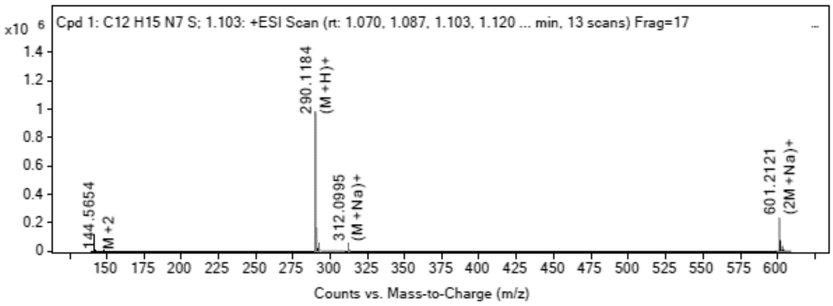

**MS Spectrum Peak List**

| Obs. *m/z* | Charge | Abund | Ion/Isotope | Tgt Mass Error (ppm) |
|---|---|---|---|---|
| 290.1184 | 1 | 986351.63 | (M+H)+ | -0.51 |
| 291.12 | 1 | 159996.56 | (M+H)+ | 1.56 |
| 292.1153 | 1 | 52298.26 | (M+H)+ | 2.25 |
| 312.0995 | 1 | 54301.61 | (M+Na)+ | 2.3 |
| 313.1017 | 1 | 8462.2 | (M+Na)+ | 2.29 |
| 314.0968 | 1 | 2829.1 | (M+Na)+ | 3.39 |
| 601.2121 | 1 | 236698.11 | (2M+Na)+ | -1.65 |
| 602.2139 | 1 | 76042.87 | (2M+Na)+ | -0.84 |
| 603.2102 | 1 | 31429.46 | (2M+Na)+ | -0.05 |
| 604.2101 | 1 | 7531.87 | (2M+Na)+ | 0.76 |

--- End Of Report ---
